# Cell-intrinsic differences between human airway epithelial cells from children and adults

**DOI:** 10.1101/2020.04.20.027144

**Authors:** Elizabeth F. Maughan, Robert E. Hynds, Adam Pennycuick, Ersilia Nigro, Kate H.C. Gowers, Celine Denais, Sandra Gómez-López, Kyren A. Lazarus, Jessica C. Orr, David R. Pearce, Sarah E. Clarke, Dani Do Hyang Lee, Maximillian N. J. Woodall, Tereza Masonou, Katie-Marie Case, Vitor H. Teixeira, Benjamin E. Hartley, Richard J. Hewitt, Chadwan Al Yaghchi, Gurpreet S. Sandhu, Martin A. Birchall, Christopher O’Callaghan, Claire M. Smith, Paolo De Coppi, Colin R. Butler, Sam M. Janes

## Abstract

The airway epithelium is a key protective barrier, the integrity of which is preserved by the self-renewal and differentiation of basal stem cells. Epithelial cells are central to the pathogenesis of multiple lung diseases. In chronic lung diseases, increasing age is a principle risk factor. Few studies have explored the differences between airway epithelial cells in children and adults and how the function of basal stem cells changes during ageing is poorly understood. Here, we analyze airway epithelial cells from children and adults in homeostatic conditions (laser capture-microdissected whole epithelium and fluorescence-activated cell-sorted basal cells) and in proliferation-inducing cell culture conditions. We find that, while the cellular composition of the pediatric and adult tracheobronchial epithelium is broadly similar, in cell culture, pediatric airway epithelial cells displayed higher colony-forming ability, sustained *in vitro* growth and outcompeted adult cells in competitive proliferation assays. In RNA sequencing experiments, we observed potentially important differences between epithelium from children and adults, including higher baseline interferon-associated gene expression in pediatric epithelium. Our results demonstrate cell-intrinsic differences in transcriptional profile and regenerative capacity between proximal airway epithelial cells of children and adults.

**Graphical abstract:** 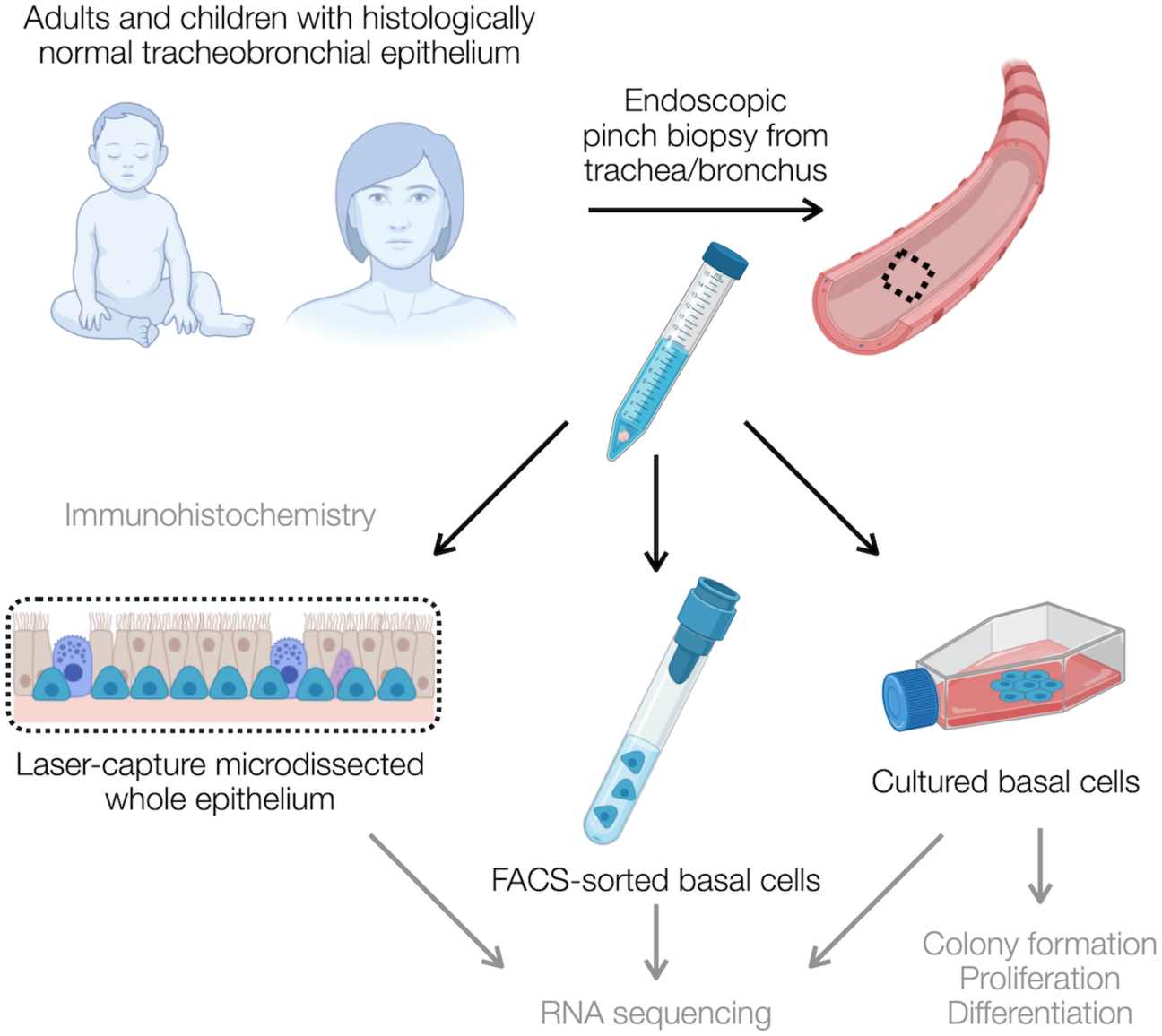

## Introduction

The human airways are lined by a pseudostratified epithelium from the trachea through most of the 16 generations of conducting airway branching. Functionally, the epithelium secretes a mucous layer and produces motile force to flow mucus proximally out of the lungs, providing protection against noxious particles and pathogens. These specialized functions are accomplished by luminal mucosecretory and ciliated epithelial cells, respectively. The differentiated cell types in this slow-turnover tissue are replenished by airway basal cells, which act as multipotent progenitors (Rock et al., 2010; Teixeira et al., 2013).

The cellular composition of the airway is being mapped in ever more detail through single cell RNA sequencing approaches, giving insight into novel cell types (Montoro et al., 2018; Plasschaert et al., 2018), differentiation intermediates in both health (Braga et al., 2019; García et al., 2019; Mori et al., 2015; Rock et al., 2011; Watson et al., 2015) and disease (Braga et al., 2019), as well as novel pathological epithelial cell subtypes, for example in idiopathic pulmonary fibrosis (Adams et al., 2020; Habermann et al., 2020; Reyfman et al., 2019). In addition to cellular diversity, region-specific differences exist in airway epithelial cell phenotype within the bronchial tree such that basal, ciliated and secretory populations in the nasal epithelium differ phenotypically (and maybe functionally) from their counterparts in the large or small airways (Braga et al., 2019; Deprez et al., 2020; Kumar et al., 2011; Travaglini et al., 2020). However, although the structural and functional consequences of ageing in the distal lung are fairly well characterized (Navarro and Driscoll, 2017; Thurlbeck and Angus, 1975; Turner et al., 1968), little is known about alterations in human proximal airway epithelial cell composition or function between pediatric and adult tissue. The importance of age-related epithelial variation has been highlighted by the COVID-19 pandemic, where functional differences in airway predisposition to viral infection and response have had a major clinical impact.

Single cell RNA sequencing studies have shown that increased transcriptional noise and upregulation of a core group of age-associated molecular pathways – including protein processing- and inflammation-associated genes – are correlated with ageing across mouse cell and tissue types, but additional processes are unique to particular cell types within specific organs, including the lungs (Angelidis et al., 2019; Kimmel et al., 2019). To date such studies have not profiled the trachea, which has distinct composition and stem cell biology to the distal lung (Basil et al., 2020), in detail. During murine tracheal ageing, epithelial cell density was reduced and the proportion of basal cells within the epithelium was slightly decreased, but there was no obvious decline in basal cell *in vitro* clonogenic potential or differentiation capacity (Wansleeben et al., 2014). However, microarray gene expression analysis of bulk tracheal cells showed changes consistent with the development of low-grade chronic inflammation in the tracheas of older mice, together with an increased presence of activated adaptive immune cells (Wansleeben et al., 2014).

There are striking differences in airway structure and composition between rodents and humans (Hogan et al., 2014), which suggests that the effects of ageing on airway epithelial regeneration might differ substantially between species. Careful characterization of human pediatric and adult airway epithelial cell composition and function will inform both lung regenerative medicine efforts and our understanding of potential pathogenic mechanisms behind multiple chronic lung disease pathologies (Kicic et al., 2006; Prasse et al., 2018; Staudt et al., 2014), as age is a risk factor for COPD, pulmonary fibrosis, infection and lung cancer. In all such diseases, basal stem cell dysfunction, perhaps accelerated by smoking, is likely to play a role in disease pathogenesis (Meiners et al., 2015).

Here, we compare homeostatic pediatric and adult human tracheobronchial epithelium in terms of cellular composition and gene expression profiles (Graphical Abstract). We further investigate proliferative airway basal stem cell behavior in primary cell culture as a surrogate for behavior during regenerative responses.

## Results

### Homeostatic human tracheobronchial epithelium has comparable cellular composition in children and adults

In mouse trachea, the proportion of the epithelium that expresses the basal cell-associated protein keratin 5 (KRT5) decreases with age (Wansleeben et al., 2014), so we first compared the cellular composition of steady state histologically normal human airway epithelium using haematoxylin and eosin (H&E) staining and immunohistochemistry for TP63 (basal cells), MUC5AC (mucosecretory cells) and FOXJ1 (ciliated cells) in tracheobronchial biopsies (Figure 1A/1B; donor characteristics are listed in Table S1). During homeostasis, we found no significant differences in the proportion of cells in these three cellular compartments in pediatric and adult biopsies either by immunohistochemistry (Figure 1A/1B), or by assessing basal, mucosecretory or ciliated cell-associated gene expression (Table S2) in bulk RNA sequencing in which we had laser capture-microdissected the whole epithelium (Figure 1C; Figure S1).

**Figure 1:**
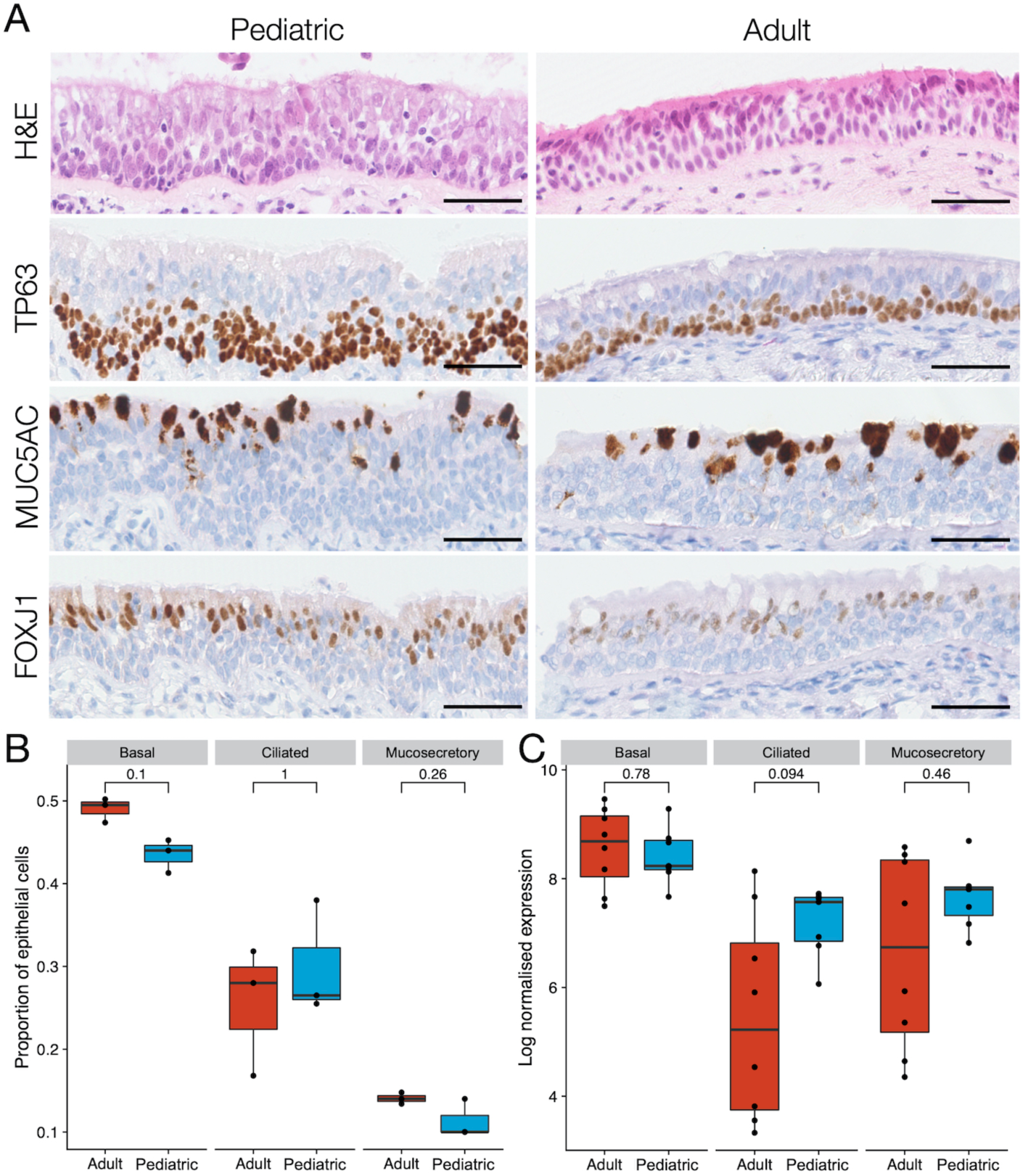
Cellular composition of pediatric and adult human tracheobronchial epithelium. **(A)** Representative haematoxylin & eosin staining and immunohistochemistry comparison of TP63, MUC5AC and FOXJ1 protein expression in pediatric and adult tracheobronchial epithelium. Scale bars = 50 μm. **(B)** Quantification of TP63^+^ basal cells, MUC5AC^+^ mucosecretory cells and FOXJ1^+^ ciliated cells in pediatric and adult tracheobronchial epithelium. Results are shown as a proportion of total cells within the epithelium (12,568 total cells for TP63, 9,651 for MUC5AC and 16,144 for FOXJ1). No significant differences were seen in a two-sided Wilcoxon rank sum test (n = 3 donors/age group; basal cells, p = 0.1; ciliated cells, p = 1; mucosecretory cells, p = 0.26). **(C)** Expression of basal, ciliated and mucosecretory cell markers in RNA sequencing data from laser capture-microdissected pediatric and adult epithelium. For each sample, the geometric mean of normalized counts of a set of cell type-specific gene markers (Table S2) is shown. No significant differences were seen in a two-sided Wilcoxon rank sum test (n = 7 pediatric and 8 adult donors; p = 0.78; ciliated cells p = 0.094, mucosecretory cells p = 0.46).

Analyzing this laser capture-microdissected whole epithelium RNA sequencing dataset using DESeq2 (Love et al., 2014) with a false discovery rate (FDR) of 1% and log_2_ fold change threshold of 1.2, we identified 72 genes with significant differential expression between pediatric and adult donors of which 40 were upregulated in adults and 32 were expressed at higher levels in children (Figure 2A; Table S3). To determine alterations in biologically functional gene groups, we performed gene set enrichment analysis (GSEA) using the Hallmark gene sets from MSigDB (Liberzon et al., 2015; Subramanian et al., 2005). In the pediatric airway epithelium, this demonstrated a higher expression of genes associated with interferon alpha and gamma responses, potentially reflecting pre-activated innate immunity in children (Loske et al., 2021). In adults, there was a higher expression of genes associated with TP53, mTORC1, Wnt-β-catenin and TGFβ signalling, as well as processes such as cholesterol homeostasis and the unfolded protein response (Figure 2B).

**Figure 2:**
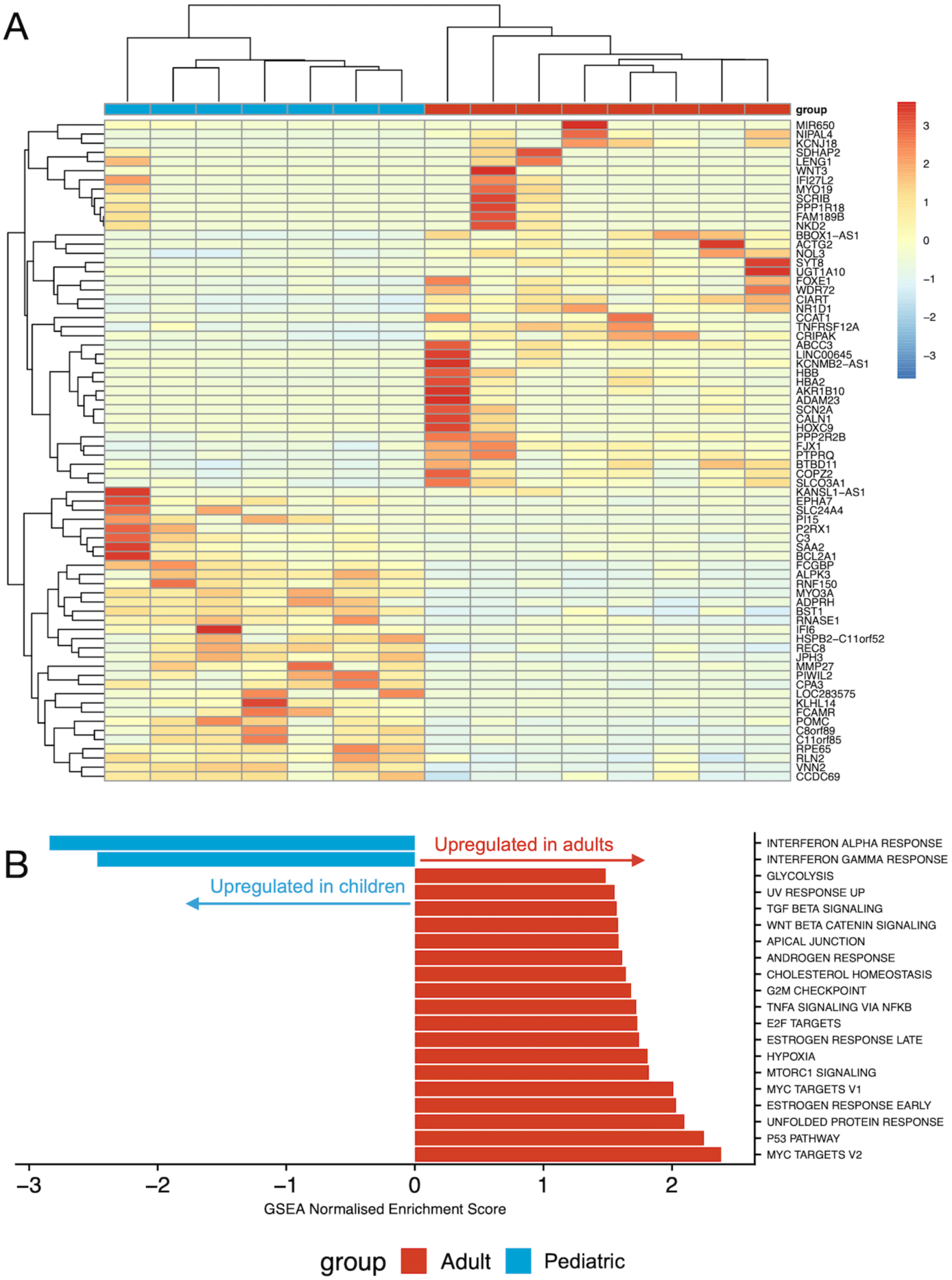
RNA sequencing of laser capture-microdissected whole epithelium from pediatric and adult proximal airways. **(A)** Cluster diagram showing the normalized expression of all 72 genes differentially expressed with a false discovery rate (FDR) < 0.01 and a log_2_ fold change > 1.2 in seven pediatric (months of age/sex; 12M, 14M, 18M, 41M, 83M, 106M) and eight adult (years of age/sex; 33F, 58F, 60F, 63M, 65M, 68F, 69M, 72M) laser capture-microdissected tracheobronchial epithelial samples. Values are scaled by row. Gene order is based on hierarchical clustering based on the similarity in overall expression patterns. Red represents relative expression higher than the median expression and blue represents lower expression. **(B)** Pathway analysis was performed on the same pediatric and adult laser capture-microdissected tracheobronchial epithelial samples using gene set enrichment analysis (GSEA) to interrogate Hallmark pathways from MSigDB. For pathways with FDR < 0.05, normalized enrichment scores are shown. A negative score (blue) represents upregulation of the pathway in the pediatric samples; a positive score (red) represents upregulation in the adult samples.

Since basal cells act as stem cells in the proximal airways, we assessed whether there are gene expression differences between pediatric and adult basal cells by using a FACS approach to isolate EpCAM^+^/PDPN^+^ basal cells (Miller et al., 2020; Weeden et al., 2017) (Figure S2) directly from tracheal biopsies. Flow cytometry experiments in independent donors verified that the majority of keratin 5 (KRT5)-expressing basal cells were included using this sorting strategy (Figure S2). Consistent with successful purification of basal cells, we saw clustering of sorted basal cells away from laser capture-microdissected whole epithelium (Figure S3A). Using DESeq2 with an FDR of 1% and log_2_ fold change threshold of 1.2, we identified 32 genes with significant differential expression between basal cells sorted from pediatric and adult donors, of which 7 were upregulated in children and 25 were more highly expressed in adults (Figure 3A; Table S3). *NTRK2* has previously been associated with basal cell function as it was upregulated in basal cells isolated from human nasal polyps compared with basal cells from the normal nasal epithelium (Ordovas-Montanes et al., 2018). Here, it was upregulated in adult compared to pediatric basal cells, consistent with a possible negative influence on basal cell progenitor function. However, the majority of differentially expressed genes do not have previously described roles in airway basal cells. GSEA suggested that pathways such as TNFα and MTORC1 signalling, as well as processes such as inflammation and apoptosis, were higher in pediatric basal cells although all pathways were of borderline statistical significance (Figure 3B).

**Figure 3:**
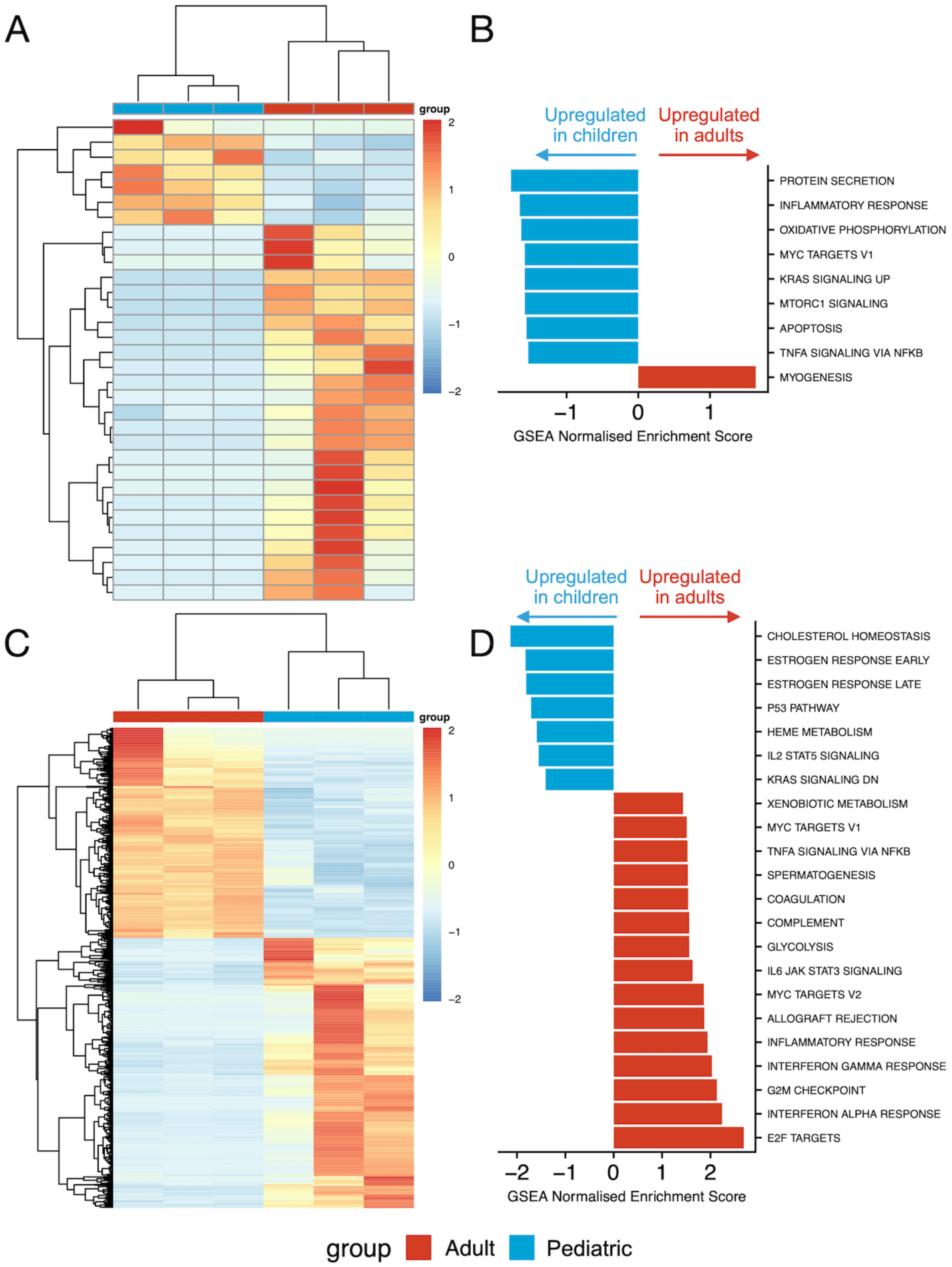
RNA sequencing of tracheobronchial basal cells from children and adults. **(A)** Cluster diagram showing the expression of 32 differentially expressed genes with a false discovery rate (FDR) < 0.01 and a log_2_ fold change > 1.2 in three pediatric (months of age/sex; 2.5M, 11M and 14M) and three adult (years of age/sex; 58F, 61F, 62F) freshly fluorescence-activated cell-sorted EpCAM^+^/PDPN^+^ basal cell samples. Gene order is based on hierarchical clustering based on the similarity in overall expression patterns. Red represents relative expression higher than the median expression and blue represents lower expression. **(B)** Pathway analysis was performed on the three pediatric and three adult freshly fluorescence-activated cell-sorted EpCAM^+^/PDPN^+^ basal cell samples using gene set enrichment analysis (GSEA) to interrogate Hallmark pathways from MSigDB. For pathways with FDR < 0.05, normalized enrichment scores are shown. A negative score (blue) represents upregulation of the pathway in the pediatric samples; a positive score (red) represents upregulation in the adult samples. **(C)** Cluster diagram showing the expression of 983 differentially expressed genes with a false discovery rate (FDR) < 0.01 and a log_2_ fold change > 1.2 in three pediatric (months of age/sex; 3M, 30M, 83M) and three adult (years of age/sex; 58F, 60F, 69M) cultured basal cell samples. Cells were cultured on 3T3-J2 mouse embryonic fibroblast feeder layers for two or three passages in epithelial cell culture medium containing Y-27632. Gene order is based on hierarchical clustering based on the similarity in overall expression patterns. Red represents relative expression higher than the median expression and blue represents lower expression. **(D)** Pathway analysis was performed on the three pediatric and three adult cultured basal cell samples using gene set enrichment analysis (GSEA) to interrogate Hallmark pathways from MSigDB. For pathways with FDR < 0.05, normalized enrichment scores are shown. A negative score (blue) represents upregulation of the pathway in the pediatric samples; a positive score (red) represents upregulation in the adult samples.

### Proliferating cultured basal cells demonstrate greater age-related transcriptional differences than basal cells *in vivo*

Given the relatively modest differences in gene expression seen in whole epithelium and FACS-sorted basal cells, we established primary cell cultures in which proliferation of airway basal cells was induced *in vitro* in order to assess whether additional differences between pediatric and adult basal cells manifest in regenerative conditions. We performed bulk RNA sequencing on cultured basal cells that were isolated and expanded on 3T3-J2 mouse embryonic feeder cells in epithelial cell culture medium containing Y-27632 (Butler et al., 2016; Hynds et al., 2019; Reynolds et al., 2016). Freshly sorted basal cells were more similar to laser capture-microdissected whole epithelium than cultured basal cells (Figure S3A), emphasizing the significant impact of the proliferation-inducing cell culture environment on the basal cell transcriptome. As expected, cultured cells were enriched for basal cell-associated genes compared to laser capture-microdissected whole epithelium (Figure S3B). Using DESeq2 with an FDR of 1% and log_2_ fold change threshold of 1.2, we identified 987 genes with significant differential expression between cultured basal cells from pediatric and adult donors, of which 555 were upregulated in children and 432 were more highly expressed in adults (Figure 3C; Table S3). GSEA suggested differences in multiple pathways between pediatric and adult cultured basal cells, some of which, including TNFα signaling and the inflammatory response, were now in the opposite direction to what was seen in basal cells *in vivo* (Figure 3D).

Notably, the mucins *MUC2, MUC3A, MUC5AC, MUC5B*, and *MUC17*, as well as the secretory master regulator *SPDEF*, were all more highly expressed by adult cultured basal cells than by pediatric basal cells. In adult basal cell cultures, we have previously observed upregulation of mucosecretory genes, such as *SCGB3A1*, in these culture conditions compared to those in a serum-free alternative, bronchial epithelial growth medium (Butler et al., 2016). However, even if mucosecretory gene expression is favored in these conditions, it is unclear why pediatric and adult epithelial cells differ in their response to culture. We hypothesized that these age-associated effects might be more pronounced after the induction of differentiation. Consistent with this, we observed that adult basal cells produced more mucous in 3D tracheosphere cultures than pediatric cells (Figure 4A; Figure S4A).

**Figure 4:**
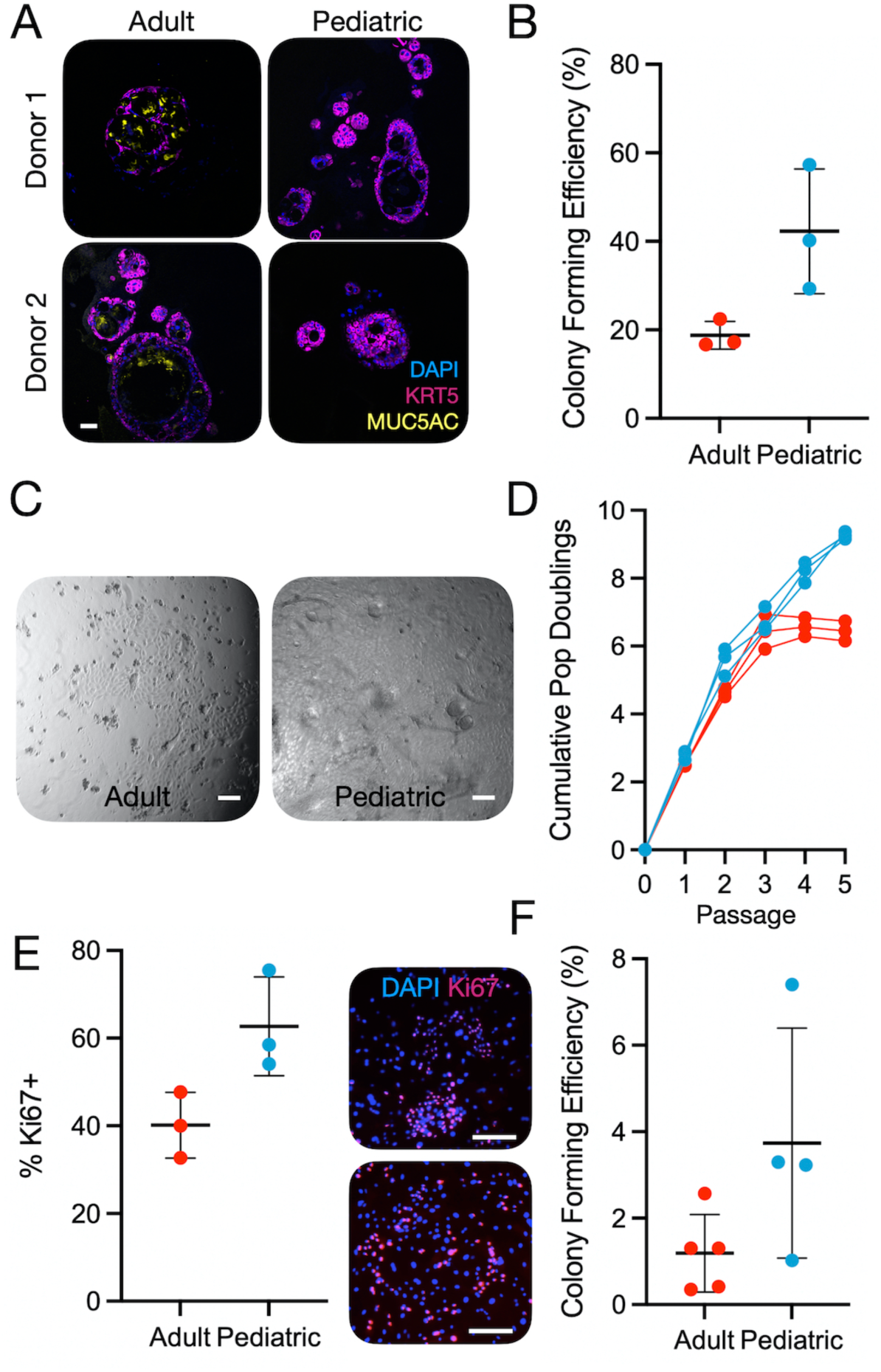
Cultured airway basal cells show cell-intrinsic differences in cell proliferation. **(A)** Immunofluorescence images showing tracheosphere cultures derived from two pediatric and two adult basal cell cultures. Sections were stained with primary antibodies against keratin 5 (KRT5; purple), MUC5AC (yellow) and a nuclear counterstain (blue). Scale bar = 50 μm. **(B)** Proportion of EpCAM^+^/PDPN^+^ basal cells that formed colonies within 7 days after sorting single cells into 96-well plates in epithelial cell culture medium containing Y-27632 (n = 3 donors/age group; n = 167-192 cells per donor; p = 0.048, two-tailed unpaired t-test). **(C)** Brightfield microscopy images showing a confluent pediatric well and an adult colony after 7 days of culture. Scale bar = 50 μm. **(D)** Population doubling analysis of the growth of pediatric and adult basal cell cultures in epithelial growth medium without Y-27632. Pediatric samples outperformed adults at passage 5 (p = 9.8×10^−5^, two-tailed unpaired t-test). **(E)** Ki67 immunofluorescence staining of passage 1 pediatric and adult basal cells cultured on feeder layers in epithelial cell culture medium without Y-27632 for 3 days before fixation. Ki67 positivity was quantified as a proportion of positive nuclei/total DAPI stained nuclei (n=3 donors/age group; mean of 1402 cells counted per donor, range 810 – 1600; p = 0.045, two-tailed unpaired t-test). The upper panel is an adult culture and the lower panel a pediatric culture. **(F)** Colony formation assays comparing early passage (P1 or P2) cultured pediatric and adult basal cells (n = 4 pediatric donors, 5 adult donors; p = 0.81, two-tailed unpaired t-test).

### Basal cells from children proliferate more readily in primary cell culture than those from adults

To investigate whether these transcriptional differences mirror functional differences between pediatric and adult basal cells, we fluorescence-activated cell-sorted single basal cells, identified by dual EpCAM (epithelial) and PDPN positivity, into individual wells of 96-well plates (range = 167 to 192 cells per donor) to compare the potential of freshly sorted pediatric and adult basal cells to generate colonies. After 7 days of culture in epithelial cell culture medium containing Y-27632, colony formation was significantly higher among basal cells derived from children than adults (Figure 4B), consistent with our previous work (Yoshida et al., 2020). Indeed, at this timepoint, pediatric basal cells had often generated colonies that had become confluent to fill the well, whereas no adult colonies reached confluence (Figure 4C; Figure S4B).

When cells were isolated and cultured in epithelial cell culture medium without Y-27632 on 3T3-J2 feeder layers, expansion of pediatric and adult cells proceeded similarly at early passages but a growth advantage was observed in pediatric donors after 5 passages (Figure 4D). In MTT assays, cultured pediatric cells showed greater proliferation than adult cells after 5 and 7 days (Figure S4C). Likewise, differences in EdU uptake (Figure S4D) and Ki67-positivity (Figure 4E) were seen in passage 1 cell cultures in the presence of feeder cells. When cultured basal cells were assessed in colony formation assays, there was a trend towards pediatric cells forming more colonies than adult cells (Figure 4F).

### Basal cells from children outcompete those from adults in mixed cultures

To better understand the differences in progenitor capacity between pediatric and adult cultures, we next developed a competitive proliferation assay (Figure S5), using lentiviral cell labelling with fluorescent constructs (Eekels et al., 2012). After optimization in 293T cells to ensure that the two lentiviruses did not affect cell growth (Figure S6), we isolated and cultured patient basal cells in epithelial cell culture medium containing Y-27632 to facilitate lentiviral transduction (Horani et al., 2013) and transduced these with either green fluorescent protein (GFP)- or mCherry-expressing lentiviral constructs (Figure 5A). When combining GFP^+^ and mCherry^+^ cells from the same donor in equal number, the ratio remained 1:1 over 7 days (Figure S7D). After transducing three pediatric and four adult basal cell cultures in this manner, we combined each pediatric donor with each adult donor in both possible color combinations so that we could monitor the growth dynamics of the two populations separately using fluorescence (Figure 5B/5C). There were no differences in lentiviral integration as determined by PCR targeting the puromycin resistance gene contained within both GFP and mCherry lentiviral vectors (Figure S7E). When cells were harvested at approximately 80% confluence, the growth differential between pediatric cells and adult cells was calculated for each pediatric/adult pair. In almost all donor pairs, the pediatric cells outgrew the adult cells (Figure 5D), although the pairings involving the youngest adult donor, who was 30 years of age, did not follow this pattern.

**Figure 5:**
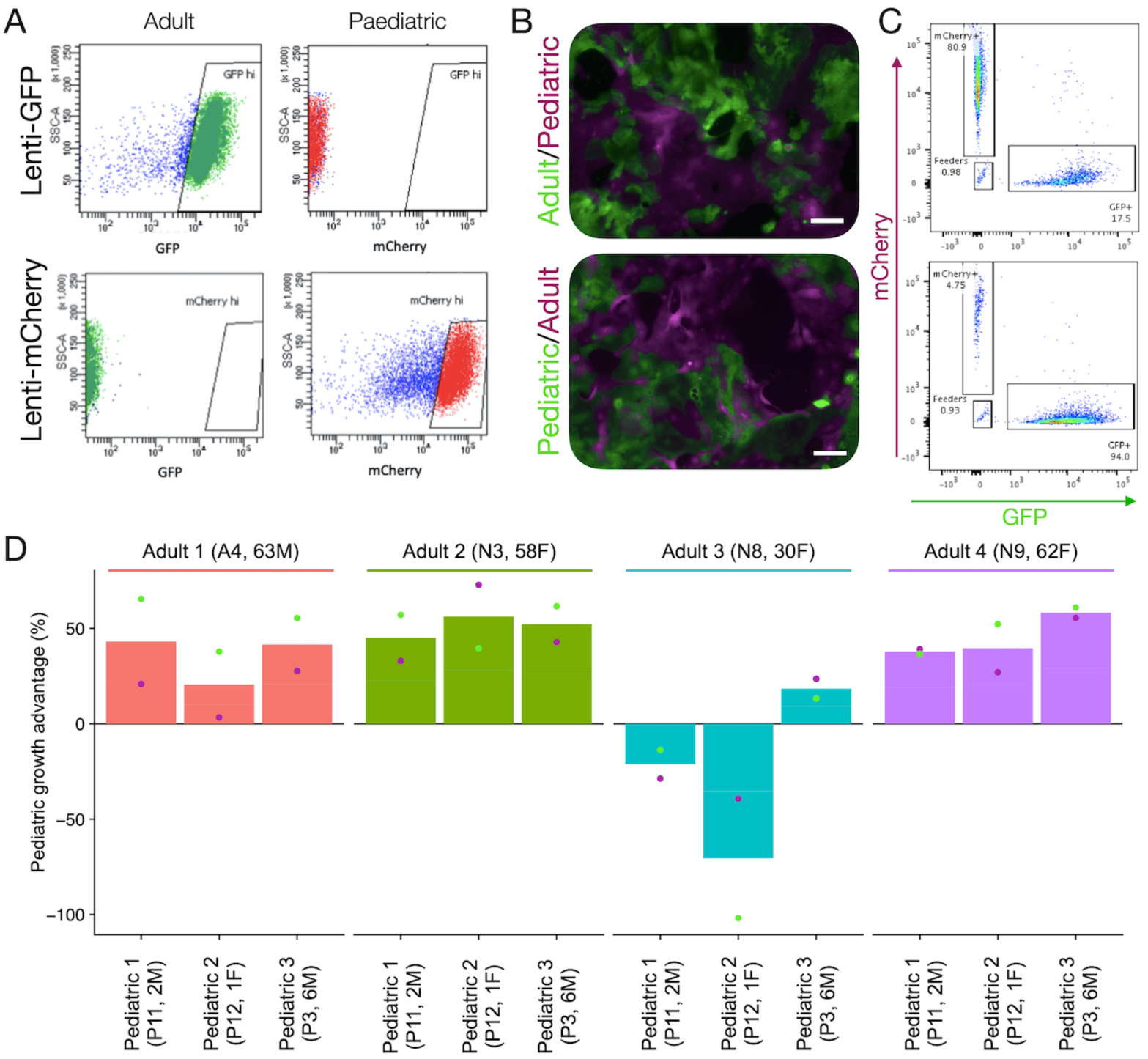
Pediatric airway basal cells out compete adult basal cells in mixed cultures. **(A)** GFP^+^ and mCherry^+^ proximal airway basal cells were generated from pediatric (n=3) and adult (n=4) donors by lentiviral transduction and purified by fluorescence-activated cell sorting (FACS). **B)** Representative immunofluorescence demonstrates how each pediatric donor was mixed with each adult donor in a 1:1 ratio by FACS in 24-well plates containing 3T3-J2 feeder cells and epithelial cell culture medium containing Y-27632. Scale bars = 200 μm. **(C)** Flow cytometric analysis was performed when cultures reached approximately 80% confluence. As shown, this allowed quantification of the abundance of GFP^+^/mCherry^+^ and GFP^-^/mCherry^-^feeder cells in these co-cultures. **(D)** Summary data for co-cultures between each pediatric and adult cell culture pair. The growth advantage of pediatric donors relative to adult donors is shown. Experiments were performed in technical triplicate and mean values for both GFP (pediatric) / mCherry (adult) and mCherry (pediatric) / GFP (adult) co-cultures are plotted as green circles and purple circles, respectively to control for possible differential effects of the specific viruses used on cell proliferation. Donor sex and age are as follows: P3 = M, 6 years old; P11 = M, 2 years old; P12 = F, 1 year old; A4 = M, 63 years old; N3 = F, 58 years old; N8 = F, 30 years old; N9 = F, 62 years old.

## Discussion

In this study, we explored differences between human tracheobronchial basal cells in children and adults in three bulk transcriptomic experiments; we compared homeostatic, laser capture-microdissected whole epithelium, homeostatic fluorescence-activated cell-sorted basal cells and proliferating cultured basal cells. At the level of the whole epithelium during homeostasis, there was broad conservation of airway epithelial transcriptional programmes although notable differences in the expression of genes associated with interferon responses were observed, consistent with a primed innate immune response in pediatric airway epithelial cells that might be associated with enhanced anti-viral response (Loske et al., 2021).

Basal cell-specific RNA sequencing revealed few differences in gene expression between pediatric and adult tracheobronchial basal cells *in vivo* that might explain the differences in their proliferative capacity once cultured, although the role of many of the differentially expressed genes has not been determined in respiratory epithelial cells. Pathway analysis suggested that, consistent with bulk epithelium, inflammatory responses are higher in pediatric basal cells suggesting that innate immune functions basal cells might also be more active in children than adults.

Upon culture, substantial transcriptomic changes were induced in airway basal cells, demonstrating one of the limitations of airway cell culture models (Orr and Hynds, 2021). However, our finding that culture conditions may affect basal cells from children differently to those from adults raises questions about whether cells from children are better able to buffer the effects of the artificial environment than those from adults. As an example, inflammatory response pathways were upregulated in adult basal cells compared to pediatric basal cells, although it has previously been shown that pediatric basal cells in these culture conditions can mount an inflammatory response upon stimulation with poly I:C (Wolf et al., 2017). These age-specific changes induced by culture should be a consideration in disease modelling using primary cells, particularly as donor age also affected the production of MUC5AC in differentiated tracheospheres.

Functional differences were observed in epithelial cell cultures where basal cells from children had a greater colony-forming capacity and proliferation than those from adults. Moreover, they had a competitive advantage in mixed cultures that was greater than would be expected from their proliferation rates as single cultures. These data are consistent with studies in other epithelia, where ageing reduces the proportion of cells identified as stem cells using *in vitro* methodologies (Barrandon and Green, 1987). Identifying whether these findings reflect *in vivo* loss of progenitor capacity with ageing will be important since this might be responsible for the differing repair responses following airway injury between children and adults (Smith et al., 2013). The molecular mechanisms by which airway basal cells lose their *in vitro* proliferative capacity with age also warrants attention, particularly given our observation that pathways associated with the inflammatory response, MYC signaling and TNF alpha signaling are higher in pediatric epithelium (and in basal cells specifically) *in vivo* but lower in adult basal cell cultures, potentially revealing a limitation of the current *in vitro* system. These data are also of relevance for lung regenerative medicine. For example, pediatric cells might be more amenable to engraftment following transplantation, as is the case in bone marrow transplantation, where donor age significantly affects outcomes (Kollman et al., 2001). Although future airway epithelial cell therapies are likely to be predominantly required by older people and be autologous in nature, recognizing age-related differences in regenerative capacity might allow the development of approaches that improve the culture and transplantation of aged basal cells. Further, corrective gene and cell therapies in the context of genetic diseases such as cystic fibrosis (Vaidyanathan et al., 2020) might be more efficient if performed early in life, when cultured cells have greater progenitor potential.

## Materials and methods

### Patient samples

Ethical approval to obtain patient tracheobronchial biopsies was granted by the National Research Ethics Committee (REC references 11/LO/1522 and 06/Q0505/12) and patients (or their parents) gave informed, written consent. Luminal biopsies were obtained using cupped biopsy forceps from patients undergoing planned rigid laryngotracheobronchoscopy under general anaesthesia or flexible bronchoscopy under sedation. Patient characteristics, procedure indication and precise site of biopsy are included in Table S1.

### Histology

Samples for histology were fixed overnight in 4% paraformaldehyde (PFA) before being dehydrated through an ethanol gradient using a Leica TP 1050 vacuum tissue processor. Samples were embedded in paraffin and sectioned at 5 µm thickness using a microtome. Haematoxylin and eosin (H&E) staining was performed using an automated system (Tissue-Tek DRS, Sakura). Immunohistochemistry was performed by The Queen Mary University of London Barts Cancer Institute Pathology Service using the Ventana DabMap Horseradish Peroxidase Kit. Stained slides were scanned using a Nanozoomer Whole Slide Imager (Hamamatsu Photonics) to create virtual slides using NDP.View2 software. For cell type quantification, images of sections that contained areas of intact epithelium with at least 300 cells were used (overall 1 – 6 slides were assessed per donor). Images were reviewed in Fiji software and positively stained cells counted using the cell count function. In total, 12,568 (TP63), 9,651 (MUC5AC) and 16,144 (FOXJ1) cells were assessed for expression of these proteins.

### Bulk RNA sequencing of tracheobronchial epithelium

Bronchoscopic biopsies were frozen immediately in optimal cutting temperature compound (OCT; in liquid hexane or on dry ice) and transported to a histopathology laboratory (Great Ormond Street Children’s Hospital, London, U.K.) within 2 hours on dry ice. Blocks and cut slides were stored at -80°C prior to use. 10 μm sections were mounted on MembraneSlide 1.0 PEN (D) membrane covered slides (Zeiss). One H&E slide was cut per block to aid navigation and identification of epithelium and basement membrane. Slides were prepared with serial washes in methanol, RNAse-free water, RNAse inhibitor and ethanol to remove residual OCT. Laser capture microdissection was performed using a PALM MicroBeam 4 Laser Microdissection microscope at 10x and 20x magnification (Figure S1) to extract the epithelial portion (or all cells above the basement membrane) from each biopsy into microadhesive-capped tubes (Zeiss). Samples were suspended in a 2:1 mix of Arcturus PicoPure Extraction buffer: RNAlater (Life Technologies) and stored at -80°C until use.

For RNA extraction, samples were thawed, disrupted by lysis (incubation at 42°C for 30 min followed by incubation at room temperature for 5 min), vortexed and filtered using RNAeasy MinElute columns (Qiagen). RNA extraction was performed using the Arcturus PicoPure RNA Isolation Kit (Life Technologies Ltd; KIT0204) as per the manufacturer’s instructions. RNA was quantified using the Qubit RNA HS Assay Kit (Thermo Fisher Scientific). Libraries were created by UCL Genomics Core Facility using the SMARTer Stranded Total RNAseq Kit (Clontech), cleaned using JetSeq (Bioline) and quality control analysis of RNA integrity was performed using High Sensitivity RNA ScreenTape and the TapeStation Analysis software (Agilent Technologies). RNA sequencing was performed using 0.5x NextSeq for 75 cycles (Illumina; 43PE, ∼33M reads per sample). RNA sequencing QC analysis is shown in Table S4.

### RNA sequencing analysis

Following sequencing, run data were demultiplexed and converted to FASTQ files using Illumina’s bcl2fastq Conversion Software v2.19. Quality control and adapter trimming were performed using fastp (Chen et al., 2018) version 0.20.1 with default settings. FASTQ files were then tagged with the UMI read (UMITools (Smith et al., 2017)) and aligned to the human genome UCSC hg38 using RNA-STAR (Dobin et al., 2013) version 2.5.2b. Aligned reads were UMI deduplicated using Je-suite (Girardot et al., 2016) version 1.2.1 and count matrices were obtained using featureCounts. Downstream analysis was performed using the R statistical environment version 3.5.0 with Bioconductor version 3.8.0 (Huber et al., 2015). Batch effects between two groups of samples run separately were identified and corrected using the *ComBat* method within the *sva* Bioconductor package (Johnson et al., 2007). Counts were compared between pediatric and adult groups using DESeq2 (Love et al., 2014) using the default settings. Where normalized counts were required, default DESeq2 normalization was applied. Pathways were assessed using an implementation of gene set enrichment analysis (GSEA) in the fgsea R package (Korotkevich et al., 2019), using Hallmark gene sets from MSigDB (Liberzon et al., 2015; Subramanian et al., 2005) as input. Heatmaps were plotted using the pheatmap (Kolde, 2012) package, implementing a complete linkage clustering method. All other plots were created using ggplot2. To avoid confounding by sex distribution, differences between the pediatric and adult groups, genes on the X and Y chromosomes were removed prior to differential analysis.

### Identification of gene lists

This study utilizes several gene lists as included in Table S2: markers of basal, secretory and ciliated cells, viral response genes and COVID-19 genes of interest. Viral genes were identified following expert review of the literature. For cell marker genes we followed the method of Danaher and colleagues (Danaher et al., 2017) to identify consistently expressed markers across age groups, based on the assumption that genes consistently associated with a cell type should correlate with each other. Candidate markers for each cell type were derived from a large single cell dataset (Travaglini et al., 2020), by taking all genes significantly over-expressed (with FDR < 0.01) in clusters associated with the given cell type. These candidate genes were then tested against healthy lung RNAseq data from the LungMAP dataset (downloaded from https://lungmap.net/breath-omics-experiment-page/?experimentTypeId=LMXT0000000018&experimentId=LMEX0000003691&analysisId=LMAN0000000342&view=allEntities; accessed on June 3, 2020), representing samples from infants, children and adults. For each list of candidate genes, we constructed a similarity matrix in each age-specific dataset following the Danaher method and performed hierarchical clustering. The final set of marker genes for each cell type were defined as those present in the main co-correlating cluster of all three age-specific datasets.

### Fluorescence-activated cell sorting of basal cells

Samples were transported to the laboratory in transport medium consisting of αMEM containing penicillin/streptomycin, amphotericin B and gentamicin. Cell suspensions were generated by sequential enzymatic digestion using dispase 16 U/mL (Corning) for 20 minutes at room temperature followed by 0.1% trypsin/EDTA (Sigma) for 30 minutes at 37°C (both in RPMI medium, Gibco). Each enzyme step was quenched with medium containing FBS, placed on ice and combined following the second digestion. Biopsies were manually homogenized using sharp dissection between digest steps and by blunt homogenization through a 100 μm cell strainer (Miltenyi Biotec). Centrifugation steps were performed at 300 x g for 5 min at 4°C.

Cells were blocked in a fluorescence-activated cell sorting (FACS) buffer composed of PBS containing 1% FBS, 25 mM HEPES buffer and 1 mM EDTA for 20 minutes in 96-well V-bottomed plates (Thermo Fisher Scientific). Cells were centrifuged as above before staining for 20 minutes on ice at 4°C. For colony formation assays and bulk RNA sequencing, the antibodies used were CD31 (BV421; Biolegend; 303124), CD45 (BV421; Biolegend; 304031), EpCAM (APC; Biolegend; 324208), PDPN (PE-Cy7; Biolegend; 337013). For flow cytometry experiments verifying that sorted populations were KRT5+ basal cells, cells were treated as above then fixed using CellFIX (BD Biosciences; 340181), permeabilized with a saponin-based reagent (Life Technologies, C10419) before intracellular staining with an anti-KRT5 antibody (AF-488; Abcam; 193894). Cells were resuspended in FACS buffer for sorting.

For colony formation assays, basal cells were sorted into collagen I-coated 96-well plates containing 3T3-J2 feeder cells at 20,000 cells per cm^2^ using a BD FACSAria Fusion FACS sorter running BD FACSDiva 8.0 software at the UCL Cancer Institute Flow Cytometry Core Facility. Experiments lasted 7 days and at termination, the number of wells that had become confluent was counted manually using a light microscope. Brightfield images were taken using a Zeiss Axiovert A1 microscope.

For bulk RNA sequencing, basal cells were sorted into epithelial cell culture medium containing Y-27632 for transport to the laboratory before being centrifuged at 300 x g for 5 min at 4°C and resuspended in RNA extraction buffer and processed as above for laser capture-microdissected samples.

### Human airway epithelial cell culture

Primary human airway epithelial cells were isolated and expanded on mitotically inactivated 3T3-J2 feeder layers in two previously reported epithelial growth media, one containing Y-27632 (Butler et al., 2016; Liu et al., 2012) and one without (Hynds et al., 2018; Rheinwald and Green, 1975). Feeder layers were prepared as previously described (Hynds et al., 2019). Epithelial cell culture medium without Y-27632 consisted of Dulbecco’s modified Eagle’s medium (DMEM)/F12 in a 3:1 ratio containing 1x penicillin–streptomycin, 10% fetal bovine serum, 1% adenine, hydrocortisone (0.5 µg/ml), EGF (10 ng/ml), insulin (5 µg/ml), 0.1 nM cholera toxin, 2 × 10^−5^ T3 and gentamicin (10 mg/ml), as previously described (Hynds et al., 2018). Epithelial cell culture medium containing Y-27632 consisted of DMEM/F12 in a 3:1 ratio containing 1x penicillin–streptomycin (Gibco), 5% fetal bovine serum (Gibco) supplemented with 5 μM Y-27632 (Cambridge Bioscience), hydrocortisone (25 ng/ml; Sigma-Aldrich), epidermal growth factor (0.125 ng/ml; Sino Biological), insulin (5 μg/ml; Sigma-Aldrich), 0.1 nM cholera toxin (Sigma-Aldrich), amphotericin B (250 ng/ml; Thermo Fisher Scientific) and gentamicin (10 μg/ml; Gibco), as previously described (Butler et al., 2016). Where indicated, dishes were collagen I-coated by diluting rat tail collagen I (BD Biosciences) to 50 μg/ml in sterile 0.02N acetic acid and applying at 5 μg/cm^2^ for one hour at room temperature in a tissue culture hood. Coated surfaces were washed once with sterile PBS before cell seeding. Population doublings were calculated as previously described (Butler et al., 2016).

RNA sequencing of cultured basal cells was performed after either two or three passages in epithelial cell culture medium containing Y-27632. RNA was extracted using the RNeasy Mini Kit (Qiagen) following the manufacturer’s instructions. RNA was submitted for library preparation by the Wellcome-MRC Cambridge Stem Cell Institute. Initial QC was performed using Qubit RNA HS Assay Kit (Thermo Fisher Scientific) and TapeStation Analysis software (Agilent Technologies). 600 ng RNA was used for library preparation using the NEBNext Ultra8482 II Directional RNA Library Prep Kit (Illumina) and the QIAseq FastSelect RNA Removal Kit (Qiagen). Libraries were subsequently measured using the Qubit system and visualized on a TapeStation D5000. Sequencing was performed using the NovaSeq6000 SP PE50 Standard (∼58M Illumina reads per sample) at the CRUK-CI Genomics Core. Following quality control and adapter trimming with fastp (as described above), paired-end reads were aligned to the human genome UCSC hg38 using RNA-STAR (Dobin et al., 2013) version 2.5.2b. Count matrices were obtained using featureCounts. Downstream analysis was performed as described for epithelial and basal RNAseq datasets above.

### Proliferation assays

For MTT assays, primary human airway epithelial cells that had been isolated and expanded in epithelial cell culture medium without Y-27632 were seeded in 96-well plates at a density of 5,000 cells/well for 24 hours without feeder cells. At the stated end-points, adherent cells were stained with MTT dye solution (10 µl of 1:10 diluted MTT stock solution in culture medium) for 3 hours at 37°C. After incubation, the medium was removed and 100 µl dimethyl sulfoxide (DMSO; Sigma-Aldrich) was added to dissolve the MTT crystals. The eluted specific stain was measured using a spectrophotometer (560 nm).

To analyze EdU uptake, passage 1 primary human airway epithelial cells that had been isolated and expanded in epithelial cell culture medium without Y-27632 were cultured until approximately 70% confluence. Cells were washed with PBS and feeder cells were removed by differential trypsinization. After washing in DMEM containing 10% FBS and then PBS again, the remaining epithelial cells were treated with 10 μM EdU (Life Technologies Click-iT EdU Alexa Fluor 488) for 1 hour. Cells were trypsinized to obtain single cell suspensions, stained according to the manufacturer’s instructions and finally co-stained with DAPI. Cells were run on an LSRFortessa (BD Biosciences) flow cytometer and data were analyzed using FlowJo 10.0.6 (TreeStar).

### Colony formation assays

2,000 cultured human airway epithelial cells were seeded per well of a six-well plate containing inactivated 3T3-J2 feeder cells. Medium was carefully changed on day 4 and day 8 of culture before the experiment was terminated on day 12. Colonies were fixed for 10 minutes in 4% PFA, stained using crystal violet (Sigma-Aldrich) at room temperature for 20 minutes and washed repeatedly in water. Colonies of more than 10 cells were counted manually using a light microscope. Colony forming efficiency was calculated as: (number of colonies formed/number of seeded cells) * 100.

### Immunofluorescence

Cells were cultured in 8-well chamber slides (Ibidi) and fixed in 4% PFA for 20 minutes at room temperature. Slides were washed and stored in PBS until staining. Cells were permeabilized and blocked in PBS containing 10% FBS and 0.025% Triton X for 1 hour at room temperature. Primary anti-Ki67 antibody (Thermo Fisher Scientific, RM-9106) was incubated overnight in block buffer without Triton X at 4°C. After three 5-minute washes in PBS, anti-rabbit secondary antibodies (AlexaFluor dyes; Molecular Probes) were incubated at a 1:200 dilution in block buffer without Triton X for 2 hours at room temperature. Cells were washed in PBS, counter-stained using DAPI (1 μg/ml stock, 1:10,000 in PBS) and washed twice more in PBS. Images were acquired using a Zeiss LSM700 confocal microscope.

Ki67 staining was performed by seeding passage 1 cells on feeder cells for 3 days. Feeder cells were removed by differential trypsinization and basal cells were fixed in 4% PFA prior to staining as above. Ki67 positivity was assessed by manual counting of Ki67-stained nuclei as a proportion of all DAPI-stained nuclei in five images per donor (mean = 1402 cells per donor, range 810 – 1600).

### GFP and mCherry lentiviral production

One Shot Stbl3 chemically competent E. coli bacteria (Thermo Fisher Scientific) were transformed using third generation lentiviral plasmids. The GFP-containing and mCherry-containing plasmids were pCDH-EF1-copGFP-T2A-Puro and pCDH-CMV-mCherry-T2A-Puro (gifts from Kazuhiro Oka; Addgene plasmids #72263 and #72264). The packaging plasmids used were pMDLg/pRRE, pRSV-Rev and pMD2.G (gifts from Didier Trono; Addgene plasmids #12251, #12253 and #12259 (Dull et al., 1998)). Bacteria were plated on ampicillin-containing agar plates overnight, and colonies expanded from this in LB broth containing ampicillin. Plasmids were extracted using the PureLinkTM HiPure Plasmid Maxiprep Kit (Thermo Fisher Scientific) with the PureLinkTM HiPure Precipitator module (Thermo Fisher Scientific) as per the manufacturer’s instructions. Plasmid DNA concentration was quantified using a NanoDrop system.

293T Human Embryonic Kidney (HEK) cells grown in DMEM containing 10% FBS and 1x penicillin/streptomycin were used to assemble and produce the viruses. Packaging and transfer plasmids were delivered into 293T cells by transfection with jetPEI (Polyplus; 101-01N) as per the manufacturer’s instructions. Medium was changed at 4 hours and 24 hours post-transfection. Culture medium containing the shed virus was collected at 72 hours and incubated with PEG-it virus precipitation solution (System Biosciences) at 4°C for 48 hours. The precipitated virus was extracted following centrifugation of the mixture as per the manufacturer’s instructions.

Viral stock titre was determined by incubating 293T HEK cells (50,000 cells/well) in 12-well plates at 1:100, 1:1000, 1:10,000 and 1:100,000 viral dilutions in complete DMEM medium containing 4 μg/mL polybrene for 4 hours before medium was refreshed. After 72 hours, cells were trypsinized, DAPI stained and GFP or mCherry positivity was determined by flow cytometry. Viral titre was calculated as:

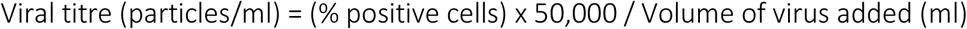

### Competitive growth assay optimization in 293T HEK cells

293T HEK cells were transduced for 24 hours with either mCherry (multiplicity of infection (MOIs) of 0.15 or 0.75) or GFP viruses (MOIs of 0.05 or 0.5). Following incubation for 72 hours, cells were sorted by FACS into high- or low-expressing populations and expanded further. Population doubling times were calculated for each color and MOI, and these values were compared to untransduced cells. Cells were then seeded as a 1:1 mix of GFP- and mCherry-expressing cells into quadruplicate wells of a 48-well plate (10,000 of each color at either high or low expression). These cells were passaged twice per week. At each passage, the contents of each well were trypsinized and the percentage of GFP and mCherry-expressing cells was analyzed by flow cytometry. 80% of each well’s cell volume was immediately re-plated into fresh 48-well plates whilst 20% of the sample from each well was stained with a live/dead fixable stain, fixed with 4% PFA and analyzed for GFP and mCherry expression by flow cytometry. Reference wells containing 10,000 cells of a single color were trypsinized at the same time points and counted manually to calculate a doubling time for each cell type.

### Lentiviral transduction of cultured human basal cells

Primary human tracheobronchial basal cells were isolated and expanded in epithelial cell culture medium containing Y-27632 for two passages before transduction with either GFP- or mCherry-containing viruses (MOI = 100). Transduction was performed in culture medium plus 4 μg/mL polybrene and medium was exchanged for fresh culture medium after 16 hours. Purification of cultures to remove untransduced cells was performed by FACS for GFP or mCherry positivity after 7-10 days of further expansion. Since both the GFP and mCherry lentiviral plasmids carry a puromycin resistance cassette, PCR copy number analysis for the puromycin cassette was performed to quantify the number of lentiviral copies incorporated per donor cell (Taqman custom copy number assay). Genomic DNA (gDNA) was extracted from each transduced cell culture using a blood & cell culture DNA mini kit (Qiagen) as per the manufacturer’s instructions. gDNA was quantified using a NanoDrop system and PCR performed using the TaqMan genotyping master mix and reference assay (Life Technologies). Stable transduction was confirmed by flow cytometry prior to competitive proliferation assays.

### Human airway basal cell competitive proliferation assays

Feeder cells were removed from cultures of GFP+ and mCherry+ basal cells by differential trypsinization and basal cells were suspended in FACS buffer containing 5 μM Y-27632 for FACS. Cells were seeded by FACS as a 1:1 mix of GFP- and mCherry-expressing cells into triplicate wells of a 24-well plate containing 3T3-J2 feeder cells (5000 or 10,000 of each color). All pediatric cultures were crossed with all of the adult cultures of the opposite label. Medium was refreshed after three and five days. At seven days or at 80-90% confluence as ascertained by fluorescence microscopy, the entire contents of the well were trypsinized for flow cytometry (i.e. feeder cells were not removed). The resulting cell suspensions were digested enzymatically with 0.25 U / mL liberase (thermolysin medium; Roche) in serum-free medium to form a single cell suspension, which was quenched with medium containing 10% FBS. Zombie Violet Live/Dead fixable stain was applied as per the manufacturer’s instructions and cell suspensions were fixed with 4% PFA for 15 min at room temperature. GFP and mCherry expression was assessed by flow cytometry. Feeder cells were seen as an unlabelled population. Untransduced pediatric cells cultured in parallel were used as non-labelled controls and single color wells containing 10,000 cells from each individual donor were used as single color controls.

Cell growth advantages were calculated using the growth calculation of Eekels *et al*. for two cell populations, assuming exponential change in the ratio of the two populations over time (Eekels et al., 2012). For example, to calculate the growth advantage of GFP-transduced cells over mCherry-transduced cells, we used the formula:

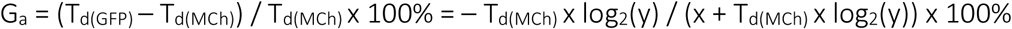

Where G_a_ is the calculated growth advantage, T_d(GFP)_ and T_d(MCh)_ are the doubling time in days of GFP+ and mCherry + cells respectively, x is the number of days of the experiment and y is equal to % GFP+ / % mCh+ at the time point x divided by the % GFP+ / % mCh+ at the time point 0. T_d(GFP)_ and T_d(MCh)_ were calculated from population doubling curves for each individual donor in these cell culture conditions. The experiment was performed in technical triplicate and for each donor pair we calculated a value for both GFP (pediatric) / mCherry (adult) and mCherry (pediatric) / GFP (adult) crosses.

## Acknowledgements

The authors thank Simon Broad and Professor Fiona Watt (Kings College London, U.K.) for providing the 3T3-J2 fibroblasts used in our study, Dr. Pascal Durrenberger (University College London, U.K.) for training and advice on laser capture microdissection, George Morrow and Dr. Barry Wilbourn (UCL Cancer Institute Flow Cytometry Core Facility, University College London, U.K.) for assistance with flow cytometry and FACS experiments, and Aimee Avery and Alex Virasami (Great Ormond Street Hospital Pathology Service, U.K.) for their assistance with cryosectioning for RNA sequencing. We also thank George Elia and his team (Pathology Service, Barts Cancer Institute, Queen Mary University of London, U.K.) for assistance with histology, Tony Brooks and Dr. Paola Niola (UCL Genomics; University College London, U.K.) for performing laser capture-microdissected whole epithelium and FACS-sorted basal cell RNA sequencing experiments and the CRUK-Cambridge Institute Genomics Core for performing cultured basal cell RNA sequencing, in particular Dr. Maike Paramor (Wellcome-MRC Cambridge Stem Cell Institute) who prepared the libraries. We further thank Samantha Arathimou (University College London), Dr. Marie-Belle El Mdawar (University College London) and Dr. Eva Grönroos (The Francis Crick Institute, U.K.) for critical reading of the manuscript.

The results published here are in part based upon data generated by the LungMAP Consortium [U01HL122642] and downloaded from (www.lungmap.net) on June 3, 2020. The LungMAP consortium and the LungMAP Data Coordinating Center (1U01HL122638) are funded by the National Heart, Lung, and Blood Institute (NHLBI).

E.F.M. (WT201265/Z/16/Z), A.P. (WT211161/Z/18/Z) and C.R.B. (WT097946MA) were Wellcome Trust Clinical Research Training Fellows. S.G.-L. was supported by a Royal Society Newton International Fellowship (NF161172). M.A.B. is an NIHR Senior Investigator (NIHR201360). P.D.C. is supported by the National Institute for Health Research (NIHR-RP-2014-04-046) and by the NIHR Great Ormond Street Hospital (GOSH) Biomedical Research Centre (BRC). R.E.H. is a Wellcome Trust Sir Henry Wellcome Fellow (WT209199/Z/17/Z) andis supported by a NIHR GOSH BRC Catalyst Fellowship. The NIHR GOSH BRC also supported the acquisition of tissue in our study. The views expressed are those of the authors and not necessarily those of the NHS, the NIHR or the Department of Health. S.M.J. was a Wellcome Trust Senior Fellow in Clinical Science (WT107963AIA). R.E.H. and S.M.J. receive funding as members of the UK Regenerative Medicine Platform (UKRMP2) Engineered Cell Environment Hub (MRC; MR/R015635/1) and P.D.C. and S.M.J. are members of the Longfonds BREATH lung regeneration consortium. R.E.H. and S.M.J. were supported by The Roy Castle Lung Cancer Foundation (2016/07/HYNDS) and S.M.J. also receives funding from The Rosetrees Trust and The UCLH Charitable Foundation.

## Author Contributions

Conceptualization, E.F.M., R.E.H., K.H.C.G., C.R.B, and S.M.J.; Methodology, E.F.M., R.E.H., K.H.C.G., C.D., S.G-L., V.H.T. and C.R.B.; Investigation, E.F.M., R.E.H., E.N., K.H.C.G., K.A.L., J.C.O., D.P., S.E.C., D.D.H.L., M.N.J.W., T.M. and K.C.; Formal Analysis, E.F.M, A.P. and R.E.H.; Resources, B.E.H., R.J.H., C.Y., G.S.S., M.A.B., C.O., C.M.S., P.D.C. and S.M.J.; Visualization, E.F.M., R.E.H. and A.P.; Writing – Original Draft, E.F.M. and R.E.H.; Writing – Review & Editing, E.F.M., R.E.H., A.P., K.H.C.G. and S.M.J.; Funding Acquisition, E.F.M., R.E.H. and S.M.J.; Supervision, R.E.H., K.H.C.G., S.G.-L, M.A.B., P.D.C., C.R.B. and S.M.J.

## Competing interests

The authors declare no competing interests relating to this manuscript. S.M.J. has attended advisory boards for Johnson and Johnson, BARD1 Life Sciences and AstraZeneca. S.M.J. receives grant funding from GRAIL Inc. and Owlstone Medical.

## Notes

### Competing Interest Statement

The authors have declared no competing interest.

### Summary of Updates

Graphical abstract added to clarify source tissue for transcriptomic datasets; additional patient samples added to Figure 2; Figure 4 revised; COVID-19 analysis of transcriptomic datasets removed given availability of larger datasets; supplemental files updated.

