## Supplementary material for "Cell-intrinsic differences between human airway epithelial cells from children and adults": Figure S

***Maughan et al.* Supplementary Data**

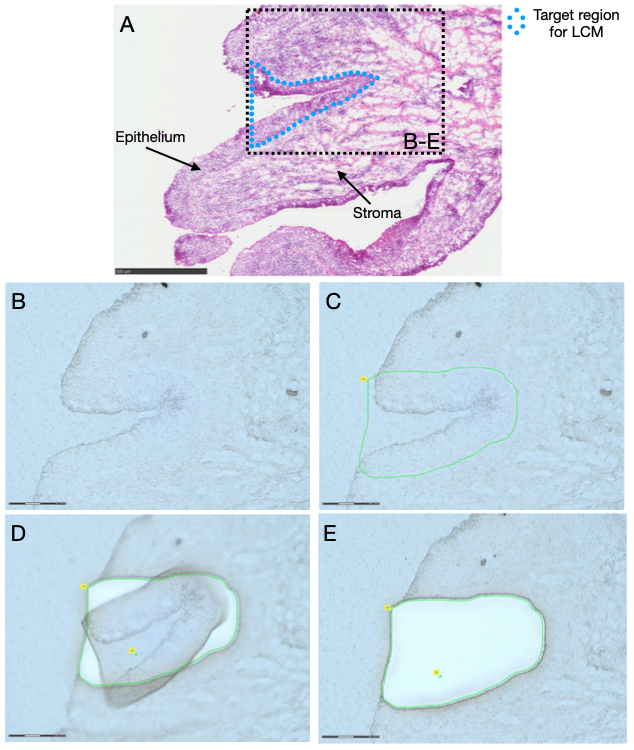

**Figure S1: Laser capture-microdissection of whole tracheobronchial epithelium from fresh frozen pediatric and adult donor tissue.**

**(A)** A slide was hematoxylin & eosin stained at the start of sectioning as a guide for identification of the epithelium. The approximate area identified in (B)-(E) is indicated by a black box. Scale bar = 500 μm.

**(B)** The equivalent area of epithelium was identified (scale bar = 150 μm) and **(C)** captured using the ‘freehand’ tool (Zeiss RoboSoftware; green line; scale bar = 150 μm). Care was taken to cut immediately below the basement membrane.

**(D)** The laser cut around this outline to release the tissue (scale bar = 150 μm).

**(E)** If samples did not immediately lift onto the adhesive surface of the collection tube cap, the ‘dot’ tool was used to capture the tissue (Zeiss RoboSoftware; scale bar = 150 μm).

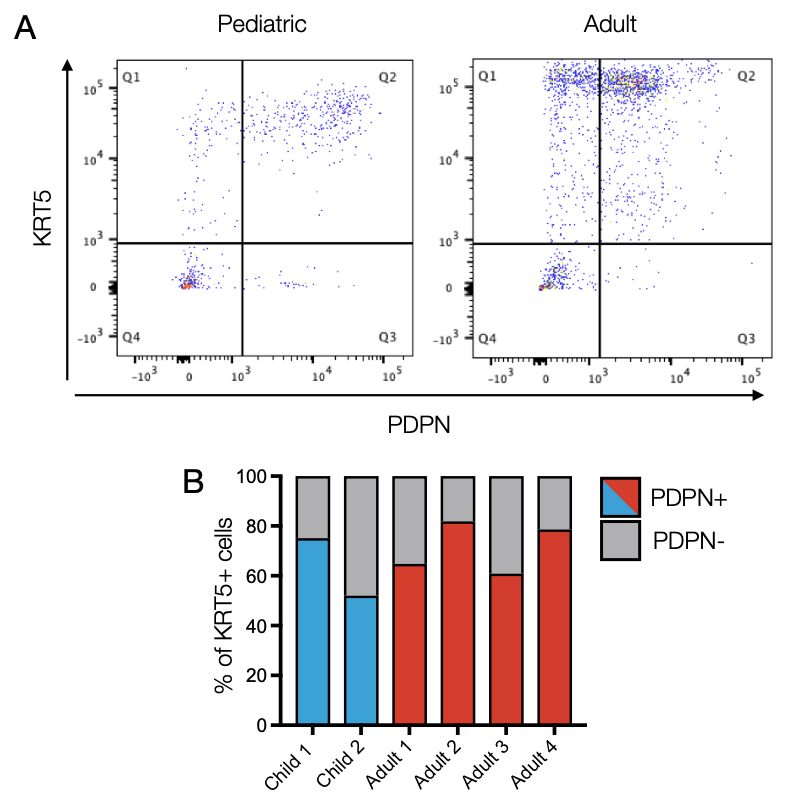

**Figure S2: Podoplanin as a human airway basal cell marker.**

**(A)** Flow cytometry analysis of podoplanin (PDPN) expression and keratin 5 expression in one pediatric (left) and one adult (right) donor. Cells were first gated as CD45-/CD31-/EpCAM+.

**(B)** Quantification of the proportion of dual KRT5+/PDPN+ cells of the total CD45-/CD31-/EpCAM+/KRT5+ population in two pediatric (blue) and four adult (red) donor tracheobronchial biopsy samples.

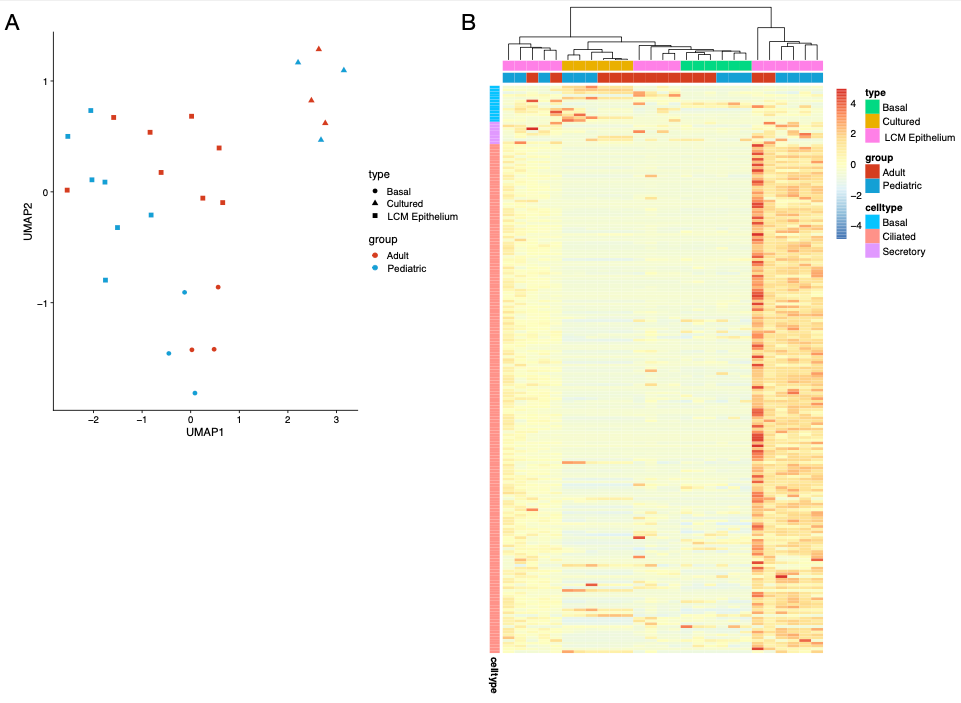

**Figure S3: Comparison of three proximal airway epithelial cell RNA sequencing datasets.**

**(A)** UMAP plot visualising laser capture-microdissected whole tracheobronchial epithelium (“LCM epithelium”, squares), fluorescence-activated cell-sorted basal cell (“basal”, circles) and cultured basal cell (“cultured”, triangles) datasets. Blue colour indicates pediatric samples and red colour indicates adult samples.

**(B)** Heatmap showing the expression of genes in our basal, mucosecretory and ciliated gene lists (see Table S2) across the three RNA sequencing datasets (laser capture-microdissected whole epithelium, “LCM epithelium”; fluorescence-activated cell-sorted EpCAM^+^/PDPN^+^ basal cells, “sorted”; cultured basal cells, “cultured”).

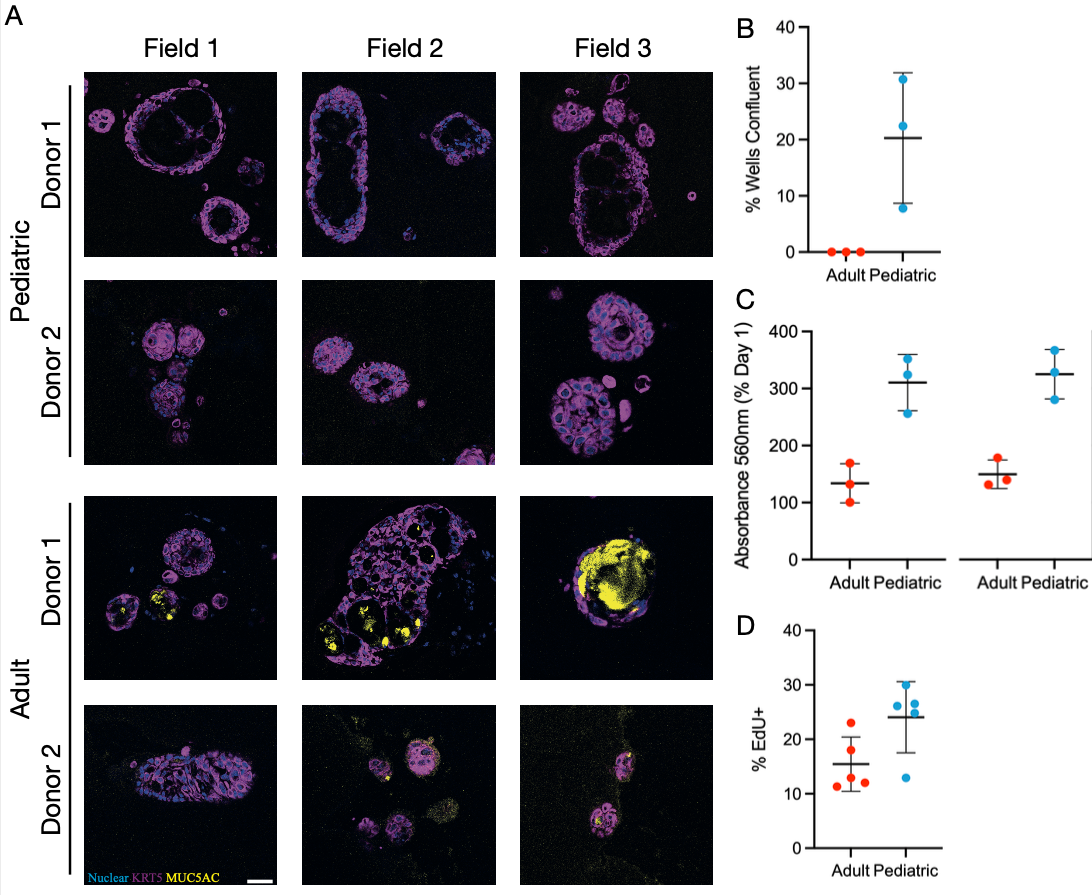
**Figure S4: Persistence of proliferation and MUC5AC-production between paediatric and adult tracheobronchial cells persist in primary cell culture.**

**(A)** Immunofluorescence imaging of tracheospheres derived from paediatric and adult cultured basal cells for keratin 5 (KRT5; purple), MUC5AC (yellow) and a nuclear counterstain (blue). Scale bar = 50 μm.

**(B)** Quantification of wells that reached confluence in 96-well plates after 14 days of culture (n = 3 donors/age group). No wells achieved confluence in the adult group.

**(C)** MTT assay assessing the growth of pediatric and adult basal cell cultures in epithelial growth medium without Y-27632 and in the absence of feeder cells. Data are normalized to readings taken at day 1. Cells were P1. 5 days (n = 3 donors/age group; p = 0.007, two-tailed unpaired t-test) and 7 days (n = 3 donors/age group; p = 0.0037, two-tailed unpaired t-test).

**(D)** Flow cytometric analysis of EdU uptake in passage 1 primary human airway epithelial cells from pediatric and adult donors grown in epithelial cell culture medium without Y-27632 at approximately 70% confluence (n = 3 donors/age group; p = 0.047, two-tailed unpaired t-test).

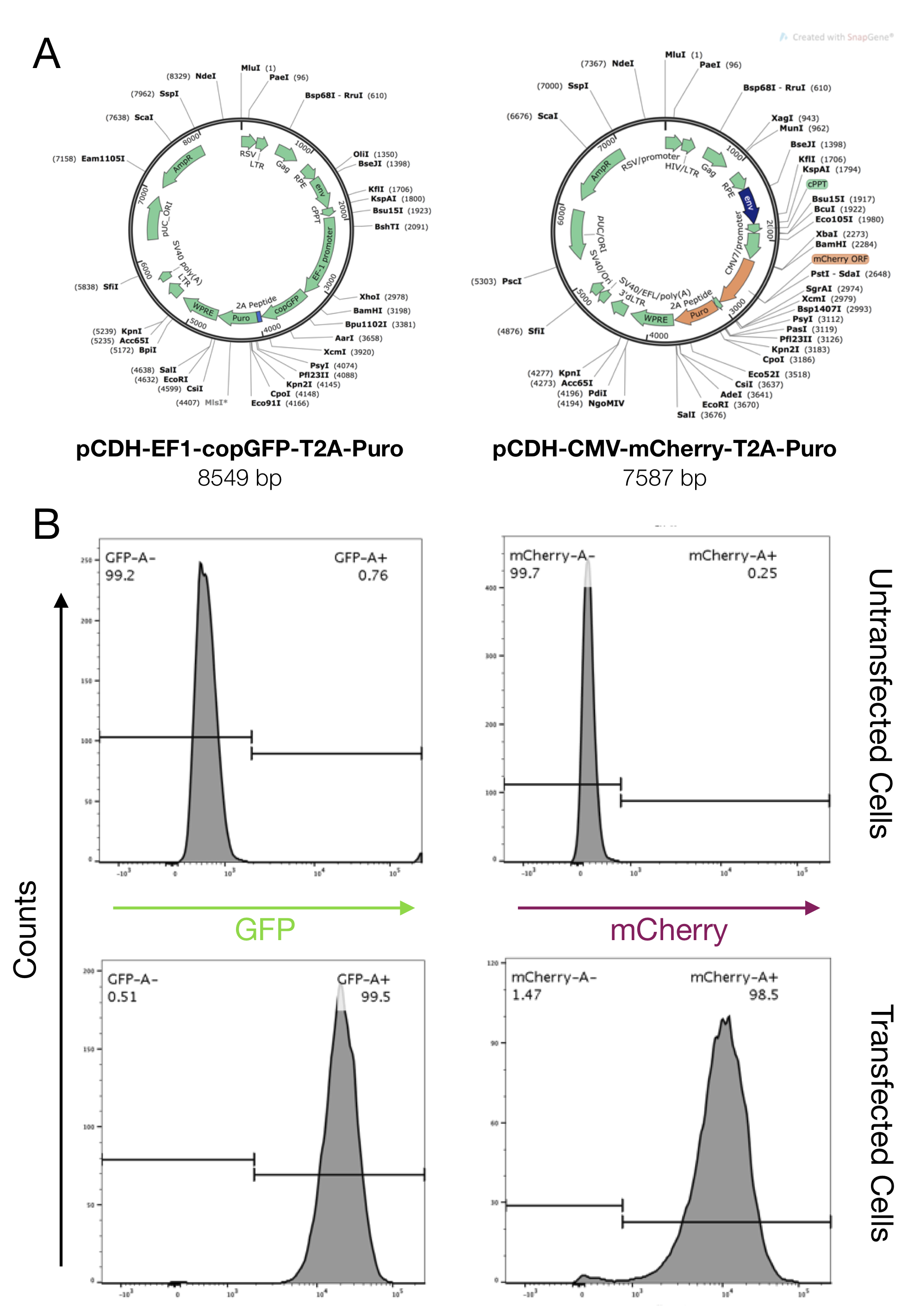

**Figure S5: Lentiviral expression of GFP or mCherry for competitive proliferation assay.**

**(A)** Plasmid maps for GFP- and mCherry-containing lentiviral plasmids (created using SnapGene software).

**(B)** Flow cytometric analysis of transfection efficiency in 293T HEK cells.

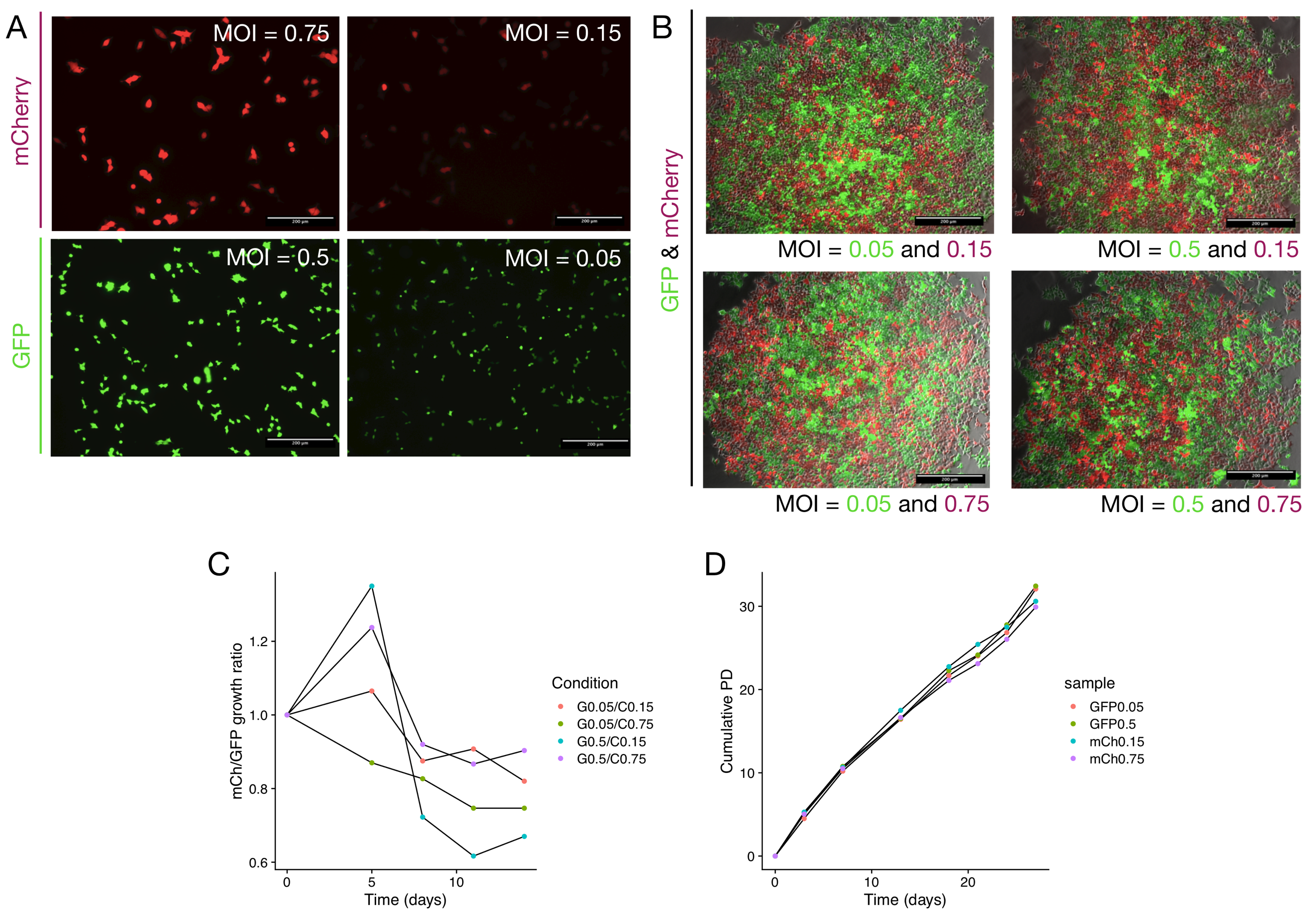

**Figure S6: Validation of competitive proliferation assay approach using 293T human embryonic kidney (HEK) cells.**

**(A)** Following lentiviral transduction at two different multiplicities of infection (MOI), 293T HEK cells were fluorescence-activated cell-sorted to generate pure cell populations of either GFP+ or mCherry+ cells. Immunofluorescence imaging showed that the fluorescence intensity was higher after transductions at the higher MOI. Cells were imaged 24 hours after sorting. Cells remained stably transduced even after transduction using the lower MOI. Scale bars = 200 μm.

**(B)** Fluorescence imaging of mixed cultures of GFP+ and mCherry+ 293T HEK cells after 5 days. Scale bars = 200 μm.

**(C)** Flow cytometry analysis of the ratio of mCherry+ and GFP+ cells in mixed cultures of transduced 293T HEK cells. Cells were initially combined in a 1:1 ratio and analyzed at the time points indicated. The cells remained close to a 1:1 ratio over two weeks of culture.

**(D)** Population doubling analysis of the growth of GFP- and mCherry-expressing 293T HEK cells.

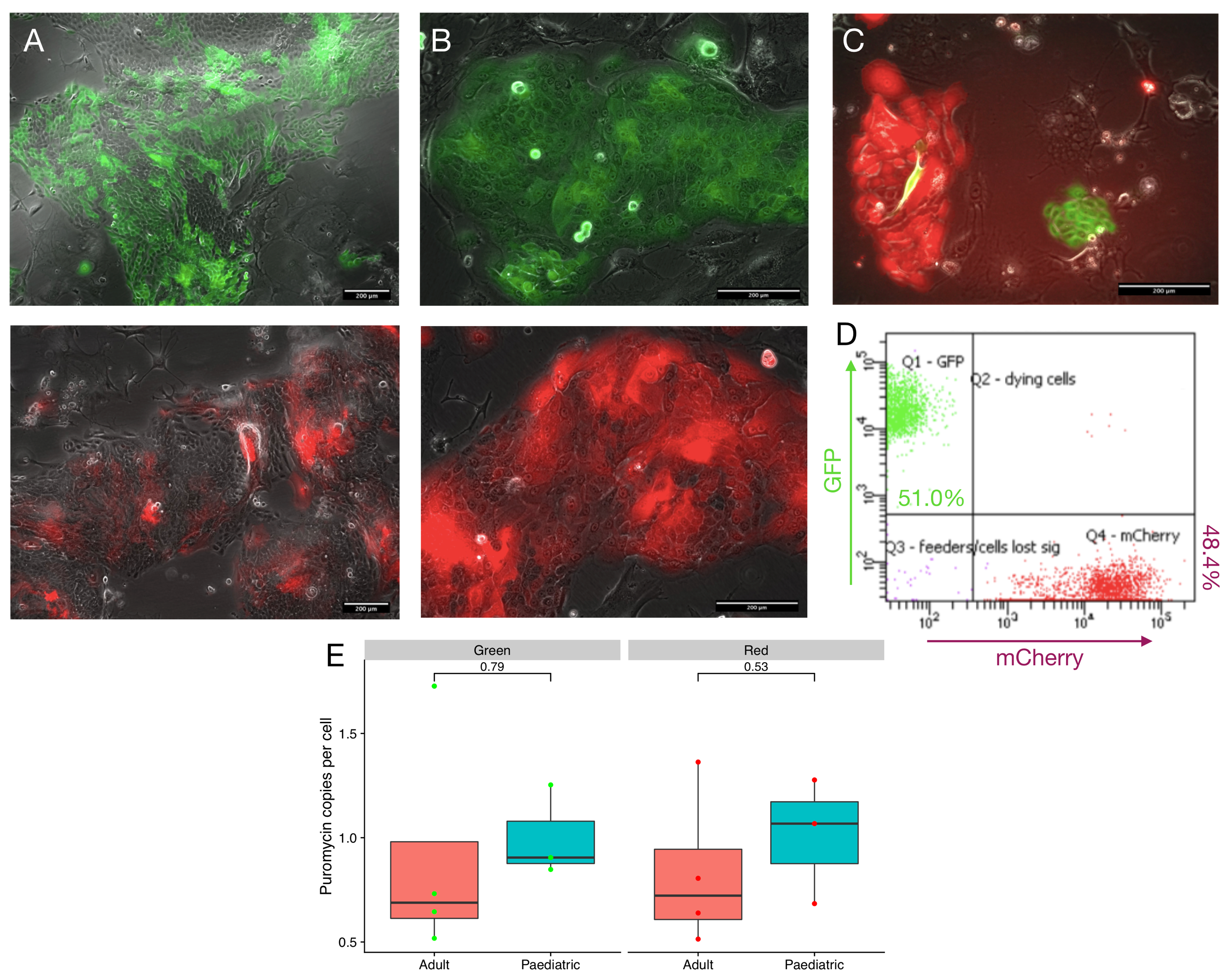

**Figure S7: Validation of competitive growth assay approach in primary human proximal airway basal cells.**

**(A)** Immunofluorescence imaging 7 days post-transduction of human basal cells with GFP (above) and mCherry (below) lentiviral constructs.

**(B)** Immunofluorescence imaging 10 days after fluorescence-activated cell sorting to purify GFP+ (above) and mCherry+ (below) populations from untransduced cells.

**(C)** Immunofluorescence imaging of cultures 4 days after mixing of primary human airway basal cells in a 1:1 ratio.

**(D)** Flow cytometry experiment in which GFP+ and mCherry+ cells from the same donor were mixed in equal ratio and analyzed after 7 days.

**(E)** Copy number determination for the puromycin resistance gene, which is present in both the GFP- and mCherry-containing lentiviruses. There were no differences in viral integration between either pediatric and adult cell cultures or between GFP and mCherry lentiviruses (p = 0.79 for GFP, p = 0.53 for mCherry, two-tailed unpaired t-test).
